# Topology of molecular deformations induces triphasic catch bonding in selectin-ligand bonds

**DOI:** 10.1101/2023.09.08.556954

**Authors:** Casey O. Barkan, Robijn F. Bruinsma

**Affiliations:** Department of Physics and Astronomy, University of California, Los Angeles, Los Angeles, CA 90095

## Abstract

Among the long-standing efforts to elucidate the physical mechanisms of protein–ligand catch bonding, particular attention has been directed at the family of selectin proteins. Selectins exhibit slip, catch-slip, and slip-catch-slip bonding, with minor structural modifications causing major changes in selectins’ response to force. How can a single structural mechanism allow interconversion between these various behaviors? We present a unifying theory of selectin-ligand catch bonding, using a structurally-motivated free energy landscape to show how the topology of force-induced deformations of the molecular system produce the full range of observed behaviors. Our novel approach can be applied broadly to other protein–ligand catch bonds, and our results have implications for such future models. In particular, our model exhibits a severe breakdown of Bell’s theory—a paradigmatic theory that is widely invoked in theories of catch bonding. This raises questions about the suitability of Bell’s theory in modeling other catch bonds.

## 1. INTRODUCTION

Mechano-sensitive proteins serve a wide range of roles in the immune system, in cell development, and cell migration [1–4]. Numerous protein-ligand bonds exhibit catch-slip bonding [5–13], where the bond’s strength varies nonmonotonically under an applied pulling force, first strengthening then weakening. Some proteins exhibit even more complex behavior: E-selectin and integrin form slip-catch-slip (triphasic) bonds with certain ligands [7, 14]. These counterintuitive responses to force have attracted much interest over the last two decades, with numerous models of catch bonding having been proposed [6, 15–23]. Yet, unresolved questions remain regarding proposed models of the selectin family of proteins (L-, P-, and E-selectin [24]). The models proposed to explain triphasic bonding either cannot account for the observed bond lifetime distributions, or appear inconsistent with selectin’s structure. Hence, it has remained an open question how triphasic bonding is produced in E-selectins, and how its mechanism relates to the mechanisms of slip and catch-slip bonding in P- and L-selectin.

The complex nature of mechanobiological systems makes modeling and model validation a challenge. Even the simplest models tend to have many fit parameters. The simple two-state catch bond model [6, 18], for example, requires eight fit parameters, and in free energy landscape models one can introduce arbitrarily many fit parameters. How can one explore, validate, and/or falsify a model while avoiding overfitting? In this work, we present a new paradigm to address this question. We develop a structural model of selectin–ligand bonding and we introduce a topological approach for classifying the mechanochemistry of the system. This topological approach allows for classification of a model’s qualitative behaviors without fine-tuning fitting parameters. With this approach, we show how selectin’s structure produces the full range of experimentally observed behaviors: slip, catch-slip, and slip-catch-slip bonding.

Our approach builds upon theoretical advances showing that catch bonding can be produced by force-induced deformation of a bond’s bound state and transition state [20, 21, 25–32].

At the heart of selectin’s distinctive mechanochemistry is an allosteric linkage between the ligand binding site and an interdomain hinge [3, 33]. While it was recognized early on that such allostery could generate catch-slip bonding [6, 34], it was not proposed that this simple mechanism could also generate slip-catch-slip behavior. Our topological approach reveals that even the simplest form of allostery contains the essential physics to produce slip-catch-slip bonding. Furthermore, this allostery provides a sensitive mechanism by which evolution can tune bond behavior. By resolving the long-standing challenge of finding a physical mechanism of slip-catch-slip bonding consistent with data on E-selectin, our model presents the first unified theory of selectin catch bonding.

Our results have broad implications for the theory of mechanochemistry. We demonstrate that Bell’s theory [35]—a paradigmatic theory invoked by influential catch bond models such as the two-pathway model, two-state model, and sliding-rebinding model—must break down in selectins. Of course, it is known theoretically that Bell’s theory, founded upon the *frozen landscape* approximation, will break down at large forces. Our model implies that even low forces cause large-scale conformational change to the transition state of the selectin–ligand bond, leading to severe violation of the frozen landscape approximation. This strong force-dependence of the transition state occurs despite the fact that the bound state is quite stiff (insensitive to force). This motivates a reexamination of other catch bonds previously modeled using Bell’s theory. Furthermore, this brings into question the suitability of steered molecular dynamics (SMD) to study protein-ligand unbinding. Though widely used to study catch bonding [19, 36–38], the artificially strong forces used in SMD may drastically alter the unbinding dynamics.

We begin by modeling the allosteric mechanism that underlies selectin’s mechanochemistry (section II), then we show how the model’s behaviors can be classified based on the topology of molecular deformations (section III). We show quantitative comparison of our model to experimental data (section IV), then discuss the inadequacy of Bell’s theory in explaining selectin’s behaviors, and discuss how our model builds upon prior work (section V).

## II. MODELING THE ALLOSTERIC MECHANISM

Selectin proteins have been heavily studied due to their importance in initiating inflammatory response [3, 24], and, as a result, extensive data is available which we use to construct and test our model. Selectin comes in three varieties: P-selectin (the first experimentally verified catch bond [5]), L-selectin, and E-selectin (which is the only one of the three that exhibits slip-catch-slip behavior [14]). Selectins extend into the blood stream from the membranes of leukocytes and of epithelial cells, where they bind ligands (chiefly PSGL-1) expressed on the surface of leukocytes traveling in the blood stream [24]. The tip of a selectin protein, where ligand binding occurs, is comprised of a lectin and an EGF-like domain connected by an interdomain hinge. Fig. 1A shows the structure of these domains (teal) bound to the ligand PSGL-1 (magenta).

**FIG. 1.**
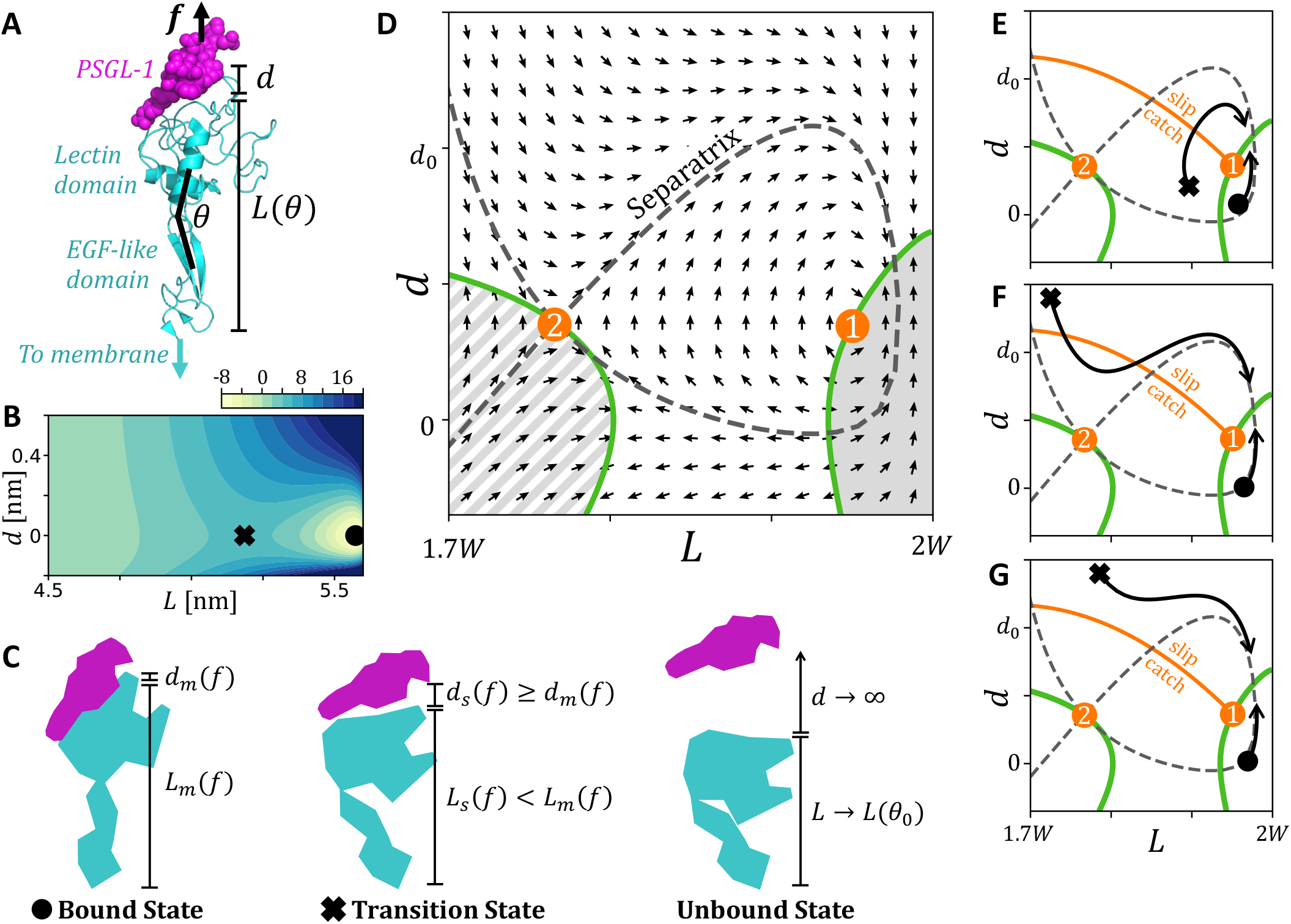
Force-induced deformation of selectin–ligand bonds. **(A)** Structure of P-selectin bound to PSGL-1 (PDB ID code: 1G1S [39], rendered in PyMOL), with interdomain hinge angle *θ*, protein extension *L*, and distance *d* between binding site and ligand. **(B)** Free energy landscape *V*_*f*_ (*L, d*) at *f* = 0 (with parameters listed in Table 1 for L-selectin). Colorbar indicates values of *V*_*f*_ in units of *k*_*B*_*T*, where 0 is the free energy of the unbound (*d → ∞*) state. The minimum point (i.e. bound state ●) and saddle point (i.e. transition state ✖) are indicated. Note that *d* = 0 corresponds to the lowest-energy bond length at zero force (hence, *d <* 0 indicates bond lengths less than this length). **(C)** Cartoon structures of selectin (teal) and ligand (magenta) in the bound state ***x***_*m*_(*f*) = (*L*_*m*_(*f*), *d*_*m*_(*f*)), transition state ***x***_*s*_(*f*) = (*L*_*s*_(*f*), *d*_*s*_(*f*)), and unbound state. **(D)** Flow of bound state and transition state under increasing *f* . det(*H*)=0 curves (green) and switch points (orange dots) are shown. Solid grey region is the minimum-like region containing the bound state. Striped grey region is the minimum-like region with no local minimum. White region is the saddle-like region containing the transition state. **(E)** Trajectories ***x***_*m*_(*f*) and ***x***_*s*_(*f*) (black curves) which cross the switch line (orange curve) from the catch region into the slip region, producing catch-slip behavior. Switch points (orange dots), switch line (orange curve), det(*H*)=0 curves (green), and separatrix (dashed grey) are shown. **(F)** Trajectories that cross the switch line from the slip region to the catch region then back to the slip region, producing slip-catch-slip behavior. **(G)** Trajectories that do not cross the switch line, producing slip behavior.

**TABLE I.** Best-fit model parameters. Units: *k*_*θ*_ (pNnm), *D*_0_ (pNnm), *d*_0_ (nm), *σ* (radians), *θ*_1_ (radians), *c* (pNnm). (*) indicates parameters that were fixed at preset values during fitting. For all selectin-ligand pairs, *W* =2.8nm and *θ*_0_ = 0.58*π* were assumed, as estimated from structure.

| Selectin | Ligand | $k_\theta$ | $D_0$ | $d_0$ | $\sigma/\pi$ | $\theta_1/\pi$ | $c$ |
| --- | --- | --- | --- | --- | --- | --- | --- |
| L-selectin | PSGL1 | 266 | 237 | 0.29 | 0.12 | 1 (*) | 0 (*) |
| LSelN138G | PSGL1 | 220 | 210 | 0.29 | 0.12 | 1 (*) | 0 (*) |
| L-selectin | 2GSP6 | 266 | 217 | 0.33 | 0.09 | 0.98 | 0 (*) |
| LSelA108H | 2GSP6 | 266 | 201 | 0.33 | 0.17 | 0.97 | 0 (*) |
| P-selectin | PSGL1 | 138 | 114 | 0.71 | 0.11 | 0.93 | 30 |
| E-selectin | sLe<sup>x</sup> | 273 | 205 | 0.26 | 0.19 | 0.99 | 10 |

An allosteric linkage between the binding site and interdomain hinge connecting lectin and EGF-like domains was revealed by experiments on mutated forms of selectin [33, 40]. These studies found that selectins bind PSGL-1 most tightly when the interdomain hinge is in an extended conformation (large *θ*, as indicated in Fig. 1A), and binding weakens as *θ* decreases. The hinge energetically prefers a bent conformation, but a pulling force exerted on the bond causes the hinge to extend, strengthening the bond and producing catch behavior. As the hinge approaches its full extension, additional force can no longer provide an additional stabilizing effect, and the bond switches to slip behavior. Hence, this simple mechanism can explain catch-slip behavior. However, can it account for slip-catch-slip behavior? We use a minimal model to investigate this question.

A minimal model for the free energy of the selectin-ligand system should retain only the essential degrees of freedom—those that are both slow *and* contribute nonharmonically to the free energy (harmonic contributions can be course-grained out, see supplemental material (SM) [41]). The allosteric coupling between the hinge and binding site implies a nonharmonic coupling between *θ* and the distance *d* between binding site and ligand, so a minimal free energy landscape should depend on *θ* and *d*. However, rather than express the free energy explicitly in terms of *θ*, it is convenient to instead use the *θ*-dependent extension of the protein, *L*, as indicated in Fig. 1A. *L* is related to *θ* by *L*(*θ*) = 2*W* sin(*θ/*2), where *W* is the length of the lectin and EGF-like domains (the two domains are nearly the same length, and we estimate *W≈* 2.8nm). Our model free energy is the following modified form of the model introduced in [32]:

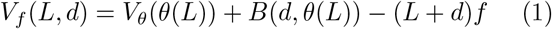

The term 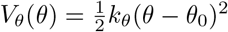 describes the hinge’s energetic preference for a bent conformation [3, 33] with preferred angle *θ*_0_ (we estimate *θ*_0_ *≈* 0.58*π* from the published structure). Binding to the ligand is described by a Morse potential *B*(*d, θ*) = *D*(*θ*)[(1 *−e*^*−d/d*^0)^2^ *−*1], where the angle-dependent binding strength *D*(*θ*) captures the allostery between hinge and binding site. The true form of *D*(*θ*) is likely complicated and presumably tuned through evolution to serve selectins’ functions. Nevertheless, a simple form for *D*(*θ*) should capture the essential physics of the system. A minimal form uses a gaussian peaked at an extended angle *θ*_1_: *D*(*θ*) = *D*_0_ exp[(*−θ −θ*_1_)^2^*/*2*σ*^2^] + *c*. This minimal form is sufficient to qualitatively explain all observed behaviors of selectins. The final term in Eq. 1, which describes the system’s coupling to the pulling force, is determined by the work done by the pulling force in bringing the system to configuration (*L, d*).

Fig. 1B plots the free energy landscape for *f* = 0. The local minimum (●), corresponding to the bound state, and saddle point (✖), corresponding to the transition state, are shown. Fig. 1C shows cartoons of the structures of the bound state, transition state, and unbound state. Importantly, these structures are force-dependent, and their force-induced deformations (section III) are key to our analysis.

To compare the model to data, predicted mean bond lifetimes *τ* must be estimated from the model. Approximate mean bond lifetime predictions can be obtained from the Arrhenius law,

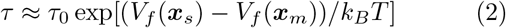

where the vectors ***x***_*m*_ = (*L*_*m*_, *d*_*m*_) and ***x***_*s*_ = (*L*_*s*_, *d*_*s*_) denote, respectively, the positions in configuration space of the local minimum (the bound state) and saddle point (the transition state) of *V*_*f*_ . The structural meaning of these quantities is illustrated in Fig. 1C. Eq. 2 is an approximation, however, and to ensure accuracy we compute *τ* by numerically solving the Fokker-Planck equation; see Appendix A for details.

Importantly, experiments measure not only mean lifetime, but also lifetime *distribution*, and models that correctly predict mean lifetime may not correctly predict the distribution of lifetimes. For a system with a single bound state, the lifetime distribution is approximately exponential: the probability that the bond ruptures at time *t* is 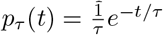 (see SM). However, in a system with multiple bound states, the lifetime distribution is *multi-*exponential (for example, FimH-mannose has two bound states and a double-exponential lifetime distribution [6]). All lifetime distribution data for selectins show a single-exponential decay [5, 14, 42], indicating only one bound state. Our proposed free energy is consistent with this experimental data, as shown by the single local minimum in Fig. 1B.

## III. TOPOLOGY OF MOLECULAR DEFORMATIONS

Eq. 2 illustrates why force-induced deformation of the bound state ***x***_*m*_ and transition state ***x***_*s*_ are key to a system’s mechanochemistry. Under increasing *f*, these deformations are described by trajectories ***x***_*m*_(*f*) and ***x***_*s*_(*f*) through the system’s configuration space, generating force-dependent bond lifetime *τ* (*f*). Topological classification of these trajectories enables qualitative predictions of a bond’s behavior without any parameter fitting.

Writing the free energy in the general form *V*_*f*_ (***x***) = *V* (***x***) *− f* ***ℓ*** · ***x*** (for Eq. 1, ***x*** = (*L, d*) and ***ℓ*** = (1, 1)), the trajectories ***x***_*m*_(*f*) and ***x***_*s*_(*f*) obey [28, 31, 32]

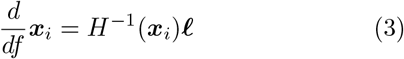

where *i* = *m* or *s* and where *H*(***x***_*i*_) is the Hessian matrix of the free energy function. A system’s catch bonding behavior is generated by the *flow* defined by this equation.

In particular, it was recently shown that force-induced switching between catch and slip behavior is generated by singularities in the flow, called *switch points* [32] (specifically, ***ℓ***-switch points, to distinguish them from ***n***-switch points; see Appendix C). The flow for our selectin model is shown in Fig. 1C, with ***ℓ***-switch points shown as orange dots labelled 1 and 2. Configuration space is divided into ‘minimum-like’ regions where minimum points can exist (where det(*H*)*>*0, shaded grey or striped in Fig. 1C), and ‘saddle-like’ regions where saddle points can exist (where det(*H*)*<*0, shaded white). Switch points lie along the boundary between these regions, where det(*H*)=0 (green curves in Fig. 1C), and a simple geometric criterion determines their location [32]. The righthand minimum-like region (solid grey) contains the bound state. The lefthand minimum-like region (striped grey) contains either no minimum point or a very shallow minimum point (see SM) for parameter values relevant to selectin.

***ℓ***-switch points shape the ‘phase portrait’ of the flow in the same way that fixed points of a dynamical system shape the system’s phase portrait. In fact, ***ℓ***-switch points are fixed points of the *regularized* flow defined by the change of variables *u* = det(*H*)*f* (see Appendix C). Under the regularized flow, switch point 1 induces circulating (locally elliptic) trajectories and switch point 2 induces locally hyperbolic trajectories. A separatrix extends from switch point 2 along the asymptotes of the locally hyperbolic flow (dashed grey curve in Fig. 1C). This separatrix separates topologically distinct families of trajectories. As we explain below, trajectories enclosed within the separatrix produce catch-slip behavior, while trajectories above the separatrix produce slip-catch-slip behavior.

The topology of the flow and of the det(*H*)=0 curve are invariant under changes in the model parameters (though very large parameter changes can modify the topology, see SM). However, the initial conditions ***x***_*m*_(0) and ***x***_*s*_(0) *do* depend on model parameters. For example, a change in parameters can shift the trajectories of the minimum (●) and saddle (✖) from those shown in Fig. 1E to those shown in Fig. 1F. The variety of observed behaviors in selectins can be explained by different initial conditions (arising due to different parameters, which reflect structural differences) that produce distinct trajectories.

The trajectories shown in Fig. 1E, 1F, and 1G illustrate how the model can exhibit catch-slip, slip-catch-slip, and slip behaviors—exactly those behaviors observed in experiments in selectins. These figures show the *switch line* (orange curve, [32]) that extends from switch point 2. The switch line indicates where the energy barrier for bond rupture switches from increasing to decreasing under increasing force. The energy barrier is increasing (catch behavior) when ***x***_*s*_ is to the bottom-left of the switch line, and the energy barrier is decreasing (slip behavior) when ***x***_*s*_ is to the top-right. Catch-slip bonding is produced by trajectories enclosed within the separatrix that cross the switch line once (Fig. 1E), and slip-catch-slip behavior is produced by trajectories above the separatrix that cross the switch line twice (Fig. 1F). For some initial conditions, the trajectory does not cross the switch line at all (Fig. 1G), producing only slip behavior. Hence, from a minimal model motivated only by available structural information, we can qualitatively explain the full range of observed behaviors while avoiding parameter fine-tuning.

## IV QUANTITATIVE COMPARISON TO DATA

While the previous section shows that the model can qualitatively capture the observed behaviors of selectin-ligand bonds, it is illuminating to quantitatively fit the model to data. Quantitative fits give insight into how trends in the experimental data relate to structural differences between different forms of selectin. Furthermore, the fits reveal how the allosteric linkage *D*(*θ*) can tune the response to force, illustrating how a wide range of behaviors may have evolved from the same simple mechanism. However, overfitting is a concern. Reassuringly, the best-fit parameters for our model are physically reasonable and the fits are obtained with between 4 and 6 free parameters (on par with or better than prior catch bond models); the method for obtaining best-fit parameters is described in the SM. Nevertheless, as with all published catch bond models, the best-fit parameters should be interpretted judiciously.

Fig. 2A shows the best-fit bond lifetime predictions for L-selectin bound to PSGL-1 (black curve), which agrees closely with experimental data (black diamonds, [43]). The best-fit parameters are given in Table 1. Fig. 2B shows the trajectories ***x***_*m*_(*f*) and ***x***_*s*_(*f*) generated using the best-fit parameters. As expected for catch-slip behavior, the trajectories are enclosed within the separatrix, and the tick mark at 64pN along the ***x***_*s*_(*f*) trajectories indicates the catch-to-slip switch. Computing predicted lifetimes requires an estimate of the friction coefficient *γ*. As discussed in Appendix A, Stokes’ law sets a lower bound of *γ* ≳ 10^*−*7^ pN s/nm, and experiments on high-friction proteins suggest an upper bound of *γ* ≲ 10^*−*2^ pN s/nm [44]. From parameter fitting, we estimate *γ ≈* 3 *×*10^*−*5^ pN s/nm, roughly halfway between these bounds.

**FIG. 2.**
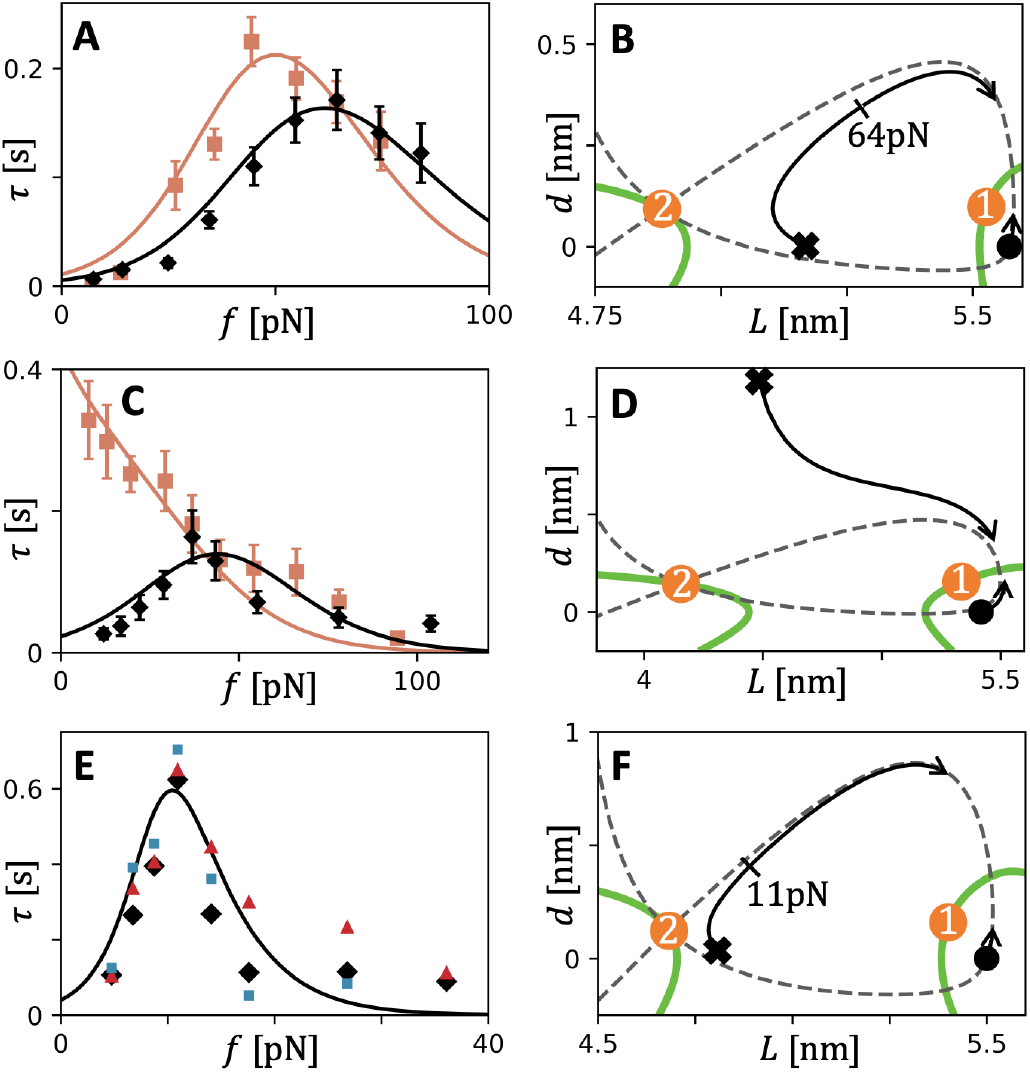
comparison of theory and data for L- and P-selectin. **(A)** L-selectin (black) and LSelN138G (orange) bound to PSGL-1. Model predictions (solid curves) and mean bond lifetime data (squares, [43]). Best fit parameters given in Table 1. **(B)** Trajectories ***x***_*s*_(*f*) and ***x***_*m*_(*f*) for L-selectin–PSGL-1, with tick mark at 64pN indicating the catch-slip switch. det(*H*)=0 curve (green), switch points (orange dots), and separatrix (grey dashed curve) are shown. **(C)** L-selectin (black) and LSelA108H (orange) bound to 2-GSP-6. Model predictions (solid curves) and mean bond life-time data (squares, [45]). This single amino acid mutation converts the catch-slip bond to a slip bond. **(D)** Trajectories ***x***_*s*_(*f*) and ***x***_*m*_(*f*) for LSelA108H–2-GSP-6. **(E)** P-selectin bound to sPSGL-1 model prediction (solid curve) and data of mean bond lifetime (black diamonds), standard deviation in bond lifetime (red triangles), and -1/slope of lifetime distribution (blue squares) [5]. **(F)** Trajectories ***x***_*s*_(*f*) and ***x***_*m*_(*f*) for P-selectin–sPSGL-1, with tick mark at 11pN indicating the catch-slip switch.

An intriguing feature of the allosteric mechanism is that the ‘spring’ of the hinge (described by *k*_*θ*_) energetically opposes ligand binding (described by *D*_0_) because binding is strongest when the hinge is extended away from its lowest-energy bent angle. This feature is reflected in the best-fit parameters: even though *k*_*θ*_ and *D*_0_ are both very large (62*k*_*B*_*T* and 55*k*_*B*_*T*, respectively), the energy barrier for bond rupture is much smaller (12*k*_*B*_*T* at *f* = 0 and 16*k*_*B*_*T* at *f* = 64pN) because the transition state finds a balance between these opposing forces.

A mutated form of L-selectin, LSelN138G, in which a key amino acid at the interdomain hinge is mutated, was found to shift the catch-slip switch to lower force and to moderately increase the maximum lifetime (Fig. 2A orange squares, [43]). Such a mutation at the interdomain hinge could be expected to modify the hinge spring constant *k*_*θ*_ and the strength of the allosteric linkage. Indeed, the close agreement between model prediction and data for LSelN138G (Fig. 2A, orange) was obtained by modifying only *k*_*θ*_ and *D*_0_ relative to the L-selectin–PSGL-1 best-fit parameters. A different mutation, within the binding site of L-selectin, converts L-selectin’s catch behavior into slip behavior when bound to the ligand 2-GSP-6 [45]. Fig. 2C shows data for L-selectin (black diamonds) and the mutated form LSelA108H (orange squares). Because this mutation is located in the binding site, we expect the hinge parameters to be unaffected by this change. Indeed, only the parameters *D*_0_, *σ*, and *θ*_1_ are modified to convert the model from L-selectin’s catch-slip behavior to LSelA108H’s slip behavior. As espected, the trajectories ***x***_*s*_(*f*) and ***x***_*m*_(*f*) corresponding to LSelA108H’s slip behavior (Fig. 2D) are topologically equivalent to the trajectories in Fig. 1G, which do not cross the switch line.

P-selectin shows a sharper catch-slip response than L-selectin, occuring over a narrower range of forces and reaching a larger maximum lifetime. Fig. 2E shows experimental bond lifetime data for P-selectin–sPSGL-1 [5]. Average measured lifetime (black squares), standard deviation of measured lifetimes (red triangles), and negative inverse slopes of measured lifetime distributions (blue squares) are shown. For an idealized bond, these three quantities would be equal, and their differences in experimental measurement roughly indicate the uncertainty in measured lifetimes [5]. A close fit to experimental data is obtained (Fig. 2E black curve, parameters in Table 1), and the trajectories ***x***_*s*_(*f*) and ***x***_*m*_(*f*) are, as expected, enclosed within the separatrix (Fig. 2F).

The slip-catch-slip behavior of E-selectin can be qualitatively described by the model in its minimal form (i.e. using the gaussian form for *D*(*θ*)), as shown in Fig. 3A (data from [14]). However, the minimal model fails to capture the narrow peak at the catch-slip switch exhibited in the data. There are many possible explanations for this discrepancy; for example, the true allosteric linkage is certainly more complicated than a simple gaussian function, and perhaps the narrow peak results from a more intricate linkage. Alternatively, perhaps the simple quadratic form that we assume for *V*_*θ*_(*θ*) is insufficient. Moreover, perhaps there are additional slow non-harmonic degrees of freedom that must be accounted for in the free energy.

**FIG. 3.**
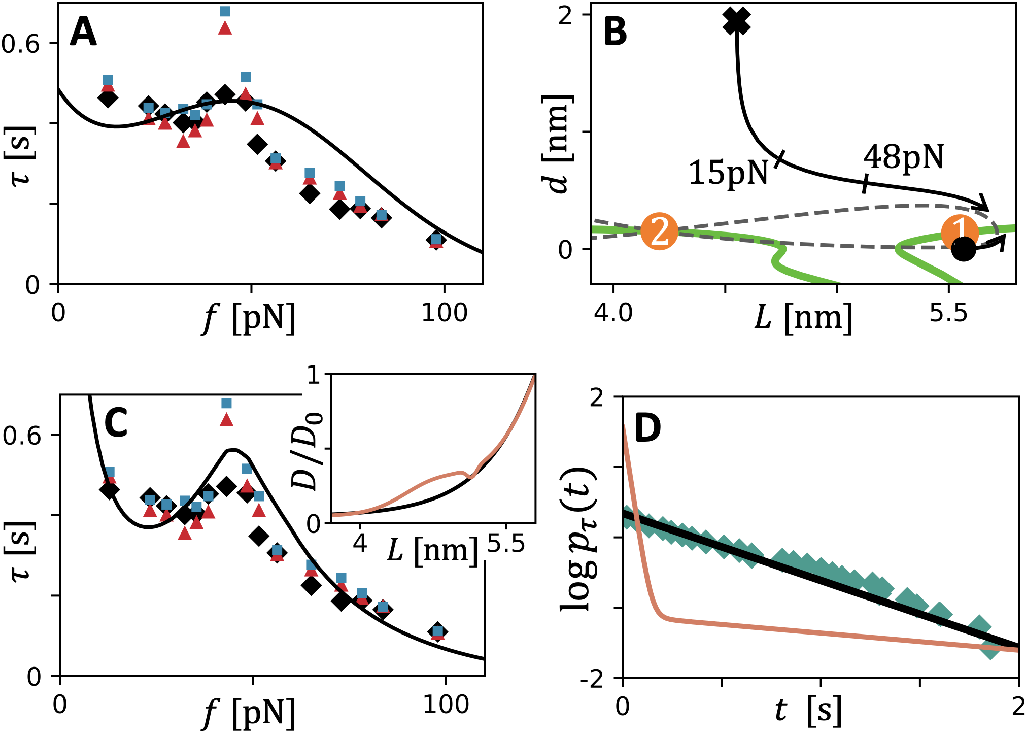
Slip-catch-slip behavior of E-selectin. **(A)** Qualitative fit to data with the minimal (gaussian) form of *D*(*θ*). Parameters given in Table 1. Data: mean bond lifetime (black diamonds), standard deviation in bond lifetime (red triangles), and -1/slope of lifetime distribution (blue squares) [14]. **(A)** Trajectories ***x***_*s*_(*f*) and ***x***_*m*_(*f*) corresponding to the theory prediction in panel A. Tick marks indicate the slip-catch switch at 15pN and catch-slip switch at 48pN. **(C)** Close agreement between theory and data achieved for the augmented allosteric linkage *D*(*θ*). Inset: augmented (orange) and gaussian (black) forms of *D*(*θ*(*L*)). The trajectories ***x***_*s*_(*f*) and ***x***_*m*_(*f*) are similar to those in panel B except that a second saddle point appears for forces between 40pN and 50pN, see SM. **(D)** Lifetime distribution for E-selectin at *f* = 48pN. Data: blue squares [14]. Our model prediction: black line. Prediction of two-state model fit to E-selectin’s slip-catch-slip data: red curve (see SM for two-state model parameters).

Although we cannot conclude definitively what causes the discrepancy between the minimal model and data for E-selectin, we can investigate whether the data can be explained by a more intricate allosteric mechanism. Specifically, we explore whether a more complex, non-gaussian, form of *D*(*θ*) can produce a closer fit to data. By augmenting *D*(*θ*) with two lower-amplitude gaussian functions, we obtain a close fit to the narrow peak in the experimental data (Fig. 3C; inset shows augmented *D*(*θ*)). The allosteric linkage in selectin proteins is presumably tuned through evolution to serve selectins’ functions, and the effect of augmenting *D*(*θ*) illustrates how the allosteric mechanism allows for evolutionary ‘tuning’ of bond behavior.

## V BELL’S THEORY AND PRIOR MODELS OF SELECTIN

Numerous models of catch bonds have been proposed in the literature, most of which involve a fixed set of bound states connected by force-dependent transition rates given by Bell’s theory [35]. Such ‘state-based’ models are defined by a set of states (both bound and ruptured states) and pathways between states, with the transition rate from state *i* to *j* via pathway *p* given by 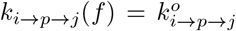. If the rates are constrained to obey detailed balance, then such a model can be viewed as a low-force approximation to a free energy landscape model, where each bound state corresponds to a local minimum of the free energy. This is known as the *frozen landscape approximation* because it neglects the movement of minimum and saddle points under the flow described by Eq. 3. In other words, the minimum and saddle points are assumed to be ‘frozen’ in place.

Our finding that ***x***_*s*_(*f*) and ***x***_*m*_(*f*) follow the nontrivial trajectories shown in Figs. 2 and 3 illustrates a failure of the frozen landscape approximation, and, therefore, a breakdown of Bell’s theory, for our model of selectin. This calls into question previously proposed state-based models of selectins and motivates a reexamination of state-based models proposed for other systems. To corroborate our claim that Bell’s theory fails for selectin, we review the prior state-based models capable of exhbiting both catch-slip and slip-catch-slip behaviors, showing how these models do not fully account for experimental data. Next, we mathematically prove that the data for E-selectin cannot be explained by any state-based model.

The most widely invoked state-based model of selectin is the *two-state model* [18, 46], a state-based model involving two connected bound states with a pathway to the ruptured state from each of the bound states. The two-state model was originally motivated by x-ray crystallography structures [39] that showed P-selectin in two conformations: an extended conformation (large *θ*) when bound to PSGL-1 and a bent conformation (small *θ*) in the absence of PSGL-1. This led to the hypothesis that the selectin–ligand complex has two meta-stable states, one bent and the other extended. The model posits that, under low force, the rupture pathway from the bent conformation is energetically favored, whereas under high force, the rupture pathway from the extended conformation is energetically favored. This contrasts with our model of selectin, where there is a single pathway (one saddle point) that continuously deforms from bent to extended under increasing force. Moreover, our model is consistent with the x-ray structures: our model predicts a bent conformation (*θ* = *θ*_0_ *≈* 0.58*π*) in the absence of PSGL-1 (i.e. in the limit that *d→ ∞*), and an extended conformation when PSGL-1 is bound (*θ ≈* 0.94*π* in the bound state shown in Fig. 1B). Experimental data of lifetime distributions can distinguish between these two models; the two-state model predicts a double-exponential *p*_*τ*_ (*t*) (Fig. 2C red curve, see SM for model parameters), whereas our model predicts a single exponential *p*_*τ*_ (*t*) (Fig. 2C black curve). Experimental data (Fig. 2C blue diamonds) is consistent with our model and falsifies the two-state model of selectin.

The sliding-rebinding model is another influential model proposed to explain both the catch-slip and slip-catch-slip behaviors of selectin-ligand bonds [14, 19]. The conceptual basis for this model involves a clever mechanism in which the pulling force exposes additional binding interactions to the ligand, strengthening the bond. This conceptual mechanism has been cast into a state-based model and fit to mean lifetime data [19]. However, this result has been criticized due to physically unrealistic fit parameters [47]. A second difficulty, which we detail in the SI, concerns the thermodynamic consistency of the state-based formulation of the model (see SI). As a result, we concur with the conclusion in [47] that this model does not capture the mechanism of selectin-ligand catch bonding.

The question remains, could there be other state-based models employing Bell’s theory that might be consistent with the data? We prove mathematically that the answer is *no* (see Appendix B). There does not exist any model based on Bell’s theory that exhibits both slip-catch-slip behavior and has only one bound state (i.e. that is consistent with mean lifetime and lifetime distribution data for E-selectin). Hence, the experiments on E-selectin show that Bell’s theory *must* break down in this system.

More recently, a free energy landscape model of selectin was proposed by Chakrabarti, Hinczewski, and Thirumalai [22] (referred to here as the CHT model), marking a major advance toward a microscopic theory of selectin’s catch bonding. This model took a mathematically elegant approach of proposing a free energy landscape for which an exact expression for mean bond lifetime was obtainable through a mean first passage time calculation. The model accurately predicts selectin’s catch-slip bond lifetime data, though the model does not exhibit slipcatch-slip behavior. The model also correctly predicts a single-exponential bond lifetime distribution, unlike the prior state-based models. While the special form of the CHT free energy was fruitful in that it allows analytical results, it also raises questions about the accuracy of the model and precludes it from exhibiting slip-catch-slip behavior. First, the model predicts that the transition state occurs at *θ* = 0 (corresponding to *θ* = *π* in the coordinates used in [22]), which would imply a non-physical overlap of the EGF-like and lectin domains. Second, the free energy landscape involves a ‘cusp’ transition state, where the bond breaks when a coordinate (*r*, in the coordinates in [22]) reaches a critical value. Modeling a free energy function with a cusp is a widely used approximation; however, the cusp removes the large *d* behavior of the free energy, which, for selectin, ‘deletes’ the portion of the free energy landscape responsible for slip-catch-slip behavior (this is apparent from the trajectory ***x***_*s*_(*f*) in Fig. 3B). In general, a cusp acts to artificially ‘pin’ the location of the saddle point (i.e. ***x***_*s*_(*f*) is pinned at one position for a range of *f*), which introduces artifacts. Our model builds on the pioneering work of the CHT model by using a free energy function that better captures the structural details. As a result, our model is the first to capture the full range of experimentally observed behaviors of selectin-ligand bonds.

## VI DISCUSSION

We propose a free energy landscape model of selectin– ligand catch bonding, the first model to explain the full range of experimentally observed bond lifetime behaviors of L-, P-, and E-selectin. By studying the topology of force-induced molecular deformations, we show how a single physical mechanism—an allosteric linkage between interdomain hinge and binding site—produces slip, catch-slip, and slip-catch-slip bonding. This topological approach allows investigation of the qualitative behaviors of the model without any parameter fitting, giving confidence that our results do not rely on finely-tuned parameters. Quantitative fits to data are also achieved with physically reasonable fitting parameters, and our model agrees with data of both mean bond lifetime and lifetime distribution. Our finding that bond behavior can be ‘tuned’ by minor changes to the allosteric linkage suggests that selectins employ a highly evolvable mechanism, letting them adapt to a variety selection pressures.

Future work applying the theory of switch points to more complex protein systems is forthcoming. T-cell receptor (TCR) binding to pMHC is one exciting system where catch bonding has been observed. Intriguingly, force-induced deformations of TCRs have been suggested to play a key role in T-cell activation [38]. However, such a model would necessitate moving beyond the 2-dimensional framework used in this work. Fortunately, the theory of switch points can be generalized to higher dimensions [32].

The violation of the frozen landscape approximation exhibited by our model, even at low forces, raises questions about the suitability of Bell’s theory for other catch bonds and other complex mechanobiological systems. Future work that develops free energy landscape models of other important catch bonds, such as integrin–fibronectin and FimH–mannose bonds, will clarify whether the frozen landscape approximation is suitable in these systems. FimH is similar to selectin in that it terminates in a lectin domain and involves an allosteric linkage at an interdomain hinge [6, 48]. Therefore, a similar model to Eq. 1 may be applicable to FimH. Importantly, FimH–mannose experiments find a double exponential lifetime distribution, necessitating a free energy model with two local minima. In fact, Eq. 1 can have two local minima (one in each minimum-like region) for certain parameter values, consistent with this data. Additionally, it has been suggested that integrin catch-slip and slip-catch-slip bonding is generated by an allosteric hinge mechanism similar to that of selectin [22]. Lifetime distributions for the integrin–fibronectin slip-catch-slip bond appear to be triple-exponential (see supplement of [7]), suggesting a more complex mechanism is at play.

It is possible that experiments could probe force-induced deformations in selectin–ligand bonds or other catch bonds, using atomic force microscopy, optical tweezers, or FRET. For selectin, our model predicts very large deformation of the transition state, yet quite small deformation of the bound state. This presents a difficulty for experiments: molecular systems pass through their transition states extremely rapidly, on far shorter timescales than the temporal resolution of experiments. The deformation of the bound state could plausibly be measured, but the predicted deformations are near or beyond the spatial resolution of current capabilities [49–51]. Other protein–ligand systems with softer bound states that undergo larger deformation may be more suited to such experiments.

An alternative to direct experimental measurement is molecular dynamics (MD) simulations. MD simulation of protein–ligand unbinding is a notoriously difficult problem [52], as typical bond lifetimes (often *∼*1s) are vastly longer than feasible simulation times (typically ≲1 ms).

To overcome this issue, steered MD simulations, in which artificially high forces are used to increase unbinding rates, are often employed to study catch bonds [19, 36– 38]. However, our results show that the transition state of catch bonds can be very sensitive to force; indeed, catch behavior is generated by this sensitivity. Hence, unbinding dynamics observed in steered MD may be drastically different from the dynamics under physiological conditions.

A recent work presents an exciting alternative to steered MD, using metadynamics to achieve the first-ever all-atom simulation of protein–ligand slip bonding under physiological forces [52]. Future refinements of this technique will likely allow all-atom simulations of catch bonding. This method requires pre-defined collective variables, and low dimensional models like our selectin model may prove very useful in substantiating choices of collective variables. More generally, low dimensional models and MD simulations should not be viewed as alternatives, but rather as complementary approaches which will likely need to be developed in tandem to provide deeper insights into molecular mechanisms of unbinding.

## Appendix A Computing mean bond lifetime from the Fokker-Planck equation

For a system with potential of mean force (i.e. free energy landscape) *V* (***x***) and uniform isotropic friction coefficient *γ*, the Fokker-Planck equation is

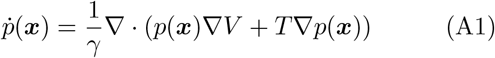

We assume such uniform isotropic friction to avoid introducing additional fit parameters. A lower bound for *γ* can be estimated from Stokes’ law (*γ* = 6*πηR* [53]) for a sphere of radius *R* in a fluid with dynamic viscosity *η*, which neglects the protein’s internal friction. For a protein in water with characteristic size of *R≈* 5nm, Stokes’ law implies *γ* ≳ 10^*−*7^ pN s/nm. A recent study on a high-friction protein system suggests *γ* ≲ 10^*−*2^ pN s/nm [44]. We estimate *γ ≈* 3 *×*10^*−*5^ pN s/nm for selectin based on fitting model prediction to data.

Mean bond lifetime can be computed from Eq. A1 by computing the quasi-stationary probability current exiting the meta-stable well. By discretizing space, this computation can be reduced to an eigenvector problem that can be numerically solved using standard linear algebra packages. We discuss the detailed implementation in the SM.

## Appendix B Proof that no state-based model can explain data for E-selectin

For E-selectin, mean bond lifetime data shows slip-catch-slip behavior and lifetime distribution data shows a single-exponential form [14]. This single-exponential form implies that there is only one bound state, and, in principle, there could be many pathways from this bound state to the ruptured state. If there are *N* rupture pathways, a state-based model employing Bell’s theory would have a rupture rate of

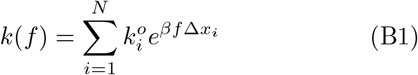

where *β* = 1*/k*_*B*_*T* and the constants ∆*x*_*i*_ may be positive or negative. Note that for *N* = 2 this is the ‘two-pathway model’ [16, 17], which was among the first theories of catch-slip bonding.

For a slip-catch-slip bond, ^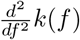^ is *negative* at the switch from slip to catch. Yet, taking the second derivative of Eq. B1 yields

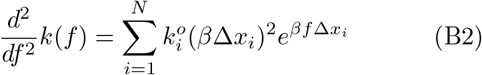

which is *always positive*. Therefore, such an *N* -pathway model cannot generate slip-catch-slip behavior and therefore cannot explain the data for E-selectin.

## Appendix C *ℓ*- and *n*-switch points, and the regularized flow equation

It was recently shown that the flow (Eq. 3) has singularities, called ***ℓ***- and ***n***-switch points, that generate force-induced switching behaviors [32]. ***ℓ***-switch points generate a switch between catch and slip via a single pathway, and they play a key role in our model of selectin. ***ℓ***-switch points are located at points on the det(*H*)=0 curve where ***v***_0_ ·***ℓ*** = 0, where ***v***_0_ is the eigenvector of *H* with vanishing eigenvalue and where ***ℓ*** is the ‘force direction’ defined above Eq. 3. ***n***-switch points are associated with a switch in pathway and occur at points on the det(*H*)=0 curve where ***v***_0_ is tangent to the det(*H*)=0 curve (i.e. where ***v***_0_ ·***n*** = 0 where ***n*** is a normal vector to the det(*H*)=0 curve). For our selectin model, there is an ***n***-switch point on the lefthand portion of the det(*H*)=0 curve shown in Fig. 1C that would indicate pathway switching from a minimum point within the lefthand minimum-like region. However, for parameter values relevant to selectin, no minimum point exists in the lefthand minimum-like region, so this ***n***-switch point has no effect on the model’s behavior.

Additionally, ***ℓ***-switch points are fixed points of the regularized flow equation obtained by the change of variables *u* = det(*H*)*f* :

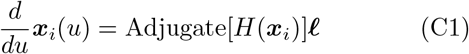

where Adjugate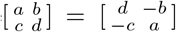 for a 2 *×* 2 matrix (in general, Adjugate[*H*] = det(*H*)*H*^*−1*^ at points where det(*H*) ≠0). This regularization removes the divergence of Eq. 3 along the det(*H*)=0 curve. The phase portrait of the regularized flow for our selectin model is shown in the SM.

