## Supplemental Material for "Topology of molecular deformations induces triphasic catch bonding in selectin-ligand bonds"

### I. SUPPLEMENTAL DISCUSSION OF THE SELECTIN–LIGAND MODEL

#### A. Model free energy (Eq. 1) with two bound states

For certain parameters, the free energy defined by Eq. 1 of the main text has two bound states (i.e. two local minima). Such a scenario is shown in Fig. SM1B. Such a free energy with two bound states may be relevant to FimH–mannose binding, for which is lifetime distribution data indicates two bound states [1].

#### B. Alternate forms of $B(d, \theta)$

In the main text, we propose a particular form for the binding free energy $B(d, \theta)$ with certain arbitrary choices. For example, we use a Morse potential to describe binding, but we could have instead used a Lennard-Jones potential. However, such changes do not modify the topology of the flow that generates slip, catch-slip, and slip-catch-slip bonding. Fig. SM1C shows the flow where the Morse potential is replaced by a Lennard-Jones potential. This change has only a minor affect on the flow.

#### C. Changing the topology with large parameter changes

As discussed in the main text, moderate changes to model parameters leave the topology of the flow and of the $\det(H)$=0 curve unchanged. However, sufficiently large changes in parameters can change the topology. This is illustrated in Fig. SM1D, where a large increase in $\sigma$ causes the two minimum-like regions to merge. Despite this change, the flow can still generate slip, catch-slip, and slip-catch-slip behaviors.

#### D. Phase portrait of regularized flow

The regularization of the flow described in Appendix C removes the divergence along the $\det(H)$=0 curve, and as a result, the direction of the flow is reversed in the saddlelike region (where $\det(H) < 0$). Fig. SM1E shows the regularized flow corresponding to Fig. 1C of the main text. For the regularized flow, the two switch points are fixed points. Switch point 1 is a fixed point with locally elliptic flow (sometimes called a 'center') and switch point 2 is a fixed point with locally hyperbolic flow (sometimes called a 'saddle', not to be confused with a saddle point of the free energy).

#### E. Trajectories for E-selectin model with augmented $D(\theta)$

The trajectories of minima and saddle points for the E-selectin model with augmented $D(\theta)$ (Fig. 3C from the main text) are shown in Fig. SM1 panels E, F, and G. The trajectories are qualitatively similar to those shown in Fig. 3B of the main text. The major difference is that for forces in the range of 49pN to 133pN, there are two saddle points. The collective effect of these two saddle points creates the sharp beak in $\tau(f)$ shown in Fig. 3C of the main text. A second apparent difference between the trajecories in Fig. SM1 and those in main text Fig. 3B is the presence of a second local minimum in Fig. SM1. However, the lefthand local minimumum in Fig. SM1 panels E-G is very shallow: $3.9 k_B T$ at $f = 0$, and lower for $f > 0$. The mean time for a system to remain in this shallow minimum before falling back into the deep (righthand) bound state) is approximately $3\mu$s, which is shorter than the mean bond lifetime by a factor of $\sim 10^6$. Hence, this shallow bound state has a negligible affect on the bond lifetime distribution.

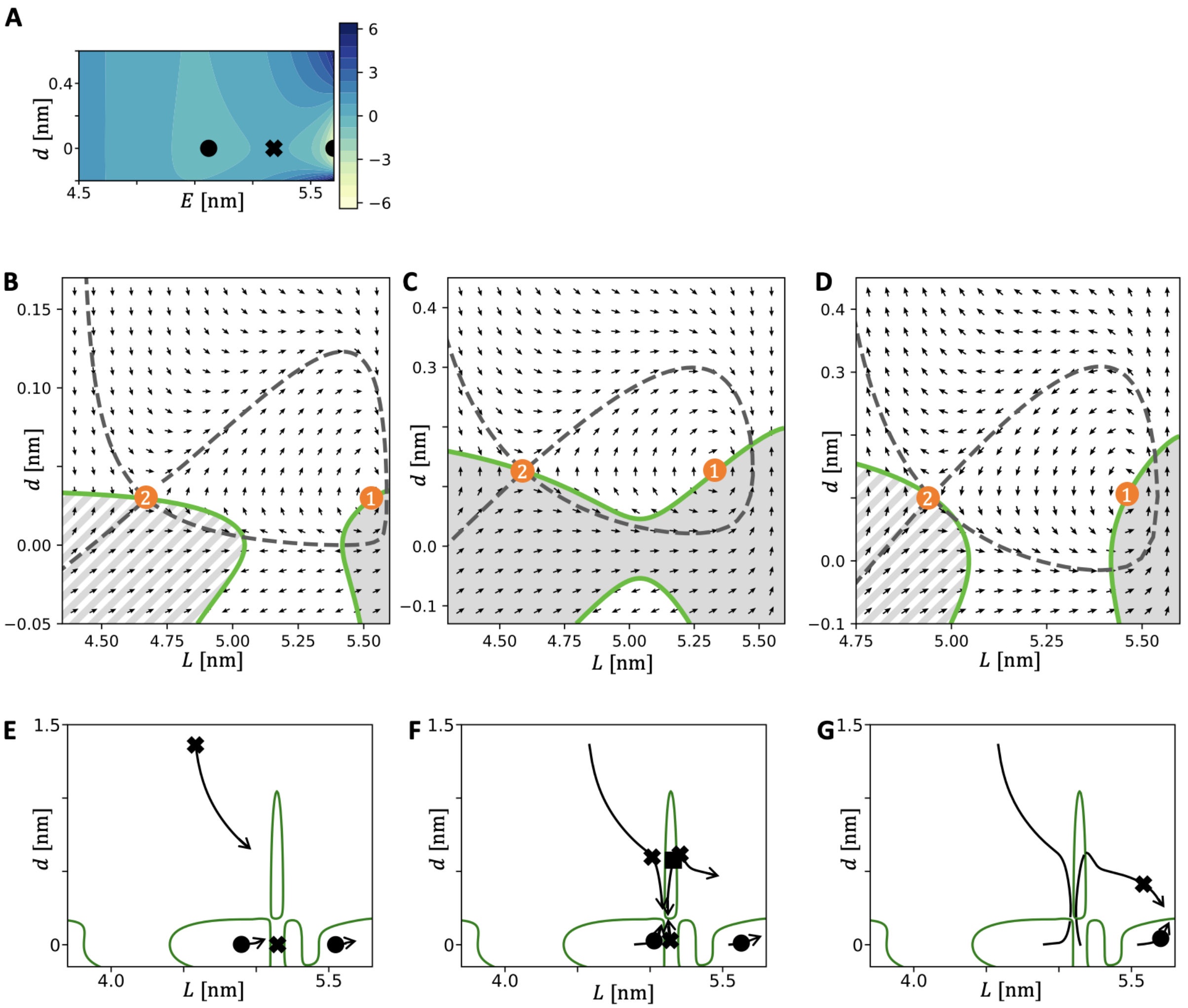


Fig. SM 1. **(A)** Free energy landscape of the form of Eq. 1 of the main text, with parameter values that produce a shallow bound state at a lower value of $E$. Colorbar shows $V_f$ at $f = 0$ in units of $k_bT$. **(B)** Flow for a free energy landscape where the Morse potential is replaced by a Lennard-Jones potential. **(C)** Flow where $\sigma$ is increased significantly, causing a change in the topology of the $\det(H)$=0 curve. The flow remains relatively unchanged. **(D)** Regularized flow, as described in Appendix C. **(E)** Trajectories of minimuma and saddle points for $f = 2$ to 48pN for the E-selectin model with augmented $D(\theta)$, as described in the main text. $\det(H)$=0 curve shown in green. **(F)** Trajectories continued up to 132pN. A maximum appears at 49pN, shown as a black square. **(G)** Trajectories continued beyond 133pN. Note that these forces are higher than any probed in experiments.

Panel E of Fig. SM1 shows the trajectories for $f = 2$pN to $f = 48$pN. Panel F shows the positions of critical points (minima, saddles, and maxima) at $f = 49$pN, and continues the trajectories up to $f = 132$pN. Panel G shows the positions of the remaining minimum and saddle at $f = 133$pN.

### F. Method of parameter fitting

The best fit parameters given in the main text were determined by first adjusting parameters by hand to obtain close fits to experimental data points, then by an automated least-squares optimization. The by-hand parameter fitting was done to ensure sensible trends in best-fit parameters between different selectin-ligand pairs (for example, the best fit $k_\theta$ for L-selectin-PSGL1 and L-selectin-2GSP6 are equal, as expected because the ligand should not affect $k_\theta$). After parameter values satisfying such constraints were found by hand, a least-squares minimization algorithm was used to collectively optimize parameters for all bonds under the specified constraints.

### II. NUMERICAL ESTIMATION OF MEAN LIFETIME USING FOKKER-PLANCK EQUATION

Under our model, the dynamics of selectin–ligand bonds are assumed to obey the Fokker-Planck (F-P) equation specified in Eq. A1 of the main text. To compute mean bond lifetime, we use the common approach of imposing boundary conditions in which bond rupture causes the system to be inserted back into the bound state. The result is that there is a steady-state solution of the F-P equation with a steady probability current from the bound state to the ruptured state. To find this steady-state current, we numerically solve the F-P equation as described below.

First, we discritize space so that (discritized) Fokker-Planck equation becomes a system of coupled ordinary differential equations of the form $\dot{\vec{p}} = M\vec{p}$ where $M$ is a square matrix of dimension equal to the number of grid points in the discritization. The null eigenvector $\vec{p}_0$ (defined by $M\vec{p}_0 = 0$) is found using the sparse eigenvector solver in the Numpy python package. The steady state current corresponding to $\vec{p}_0$ is computed, which gives the rate of bond rupture ($1/\tau$).

### III. MULTI-EXPONENTIAL LIFETIME DISTRIBUTIONS IN MULTI-STATE MODELS

For a state-based model with $M$ bound states, the bond lifetime distribution $p_\tau(t)$ is given by a sum of $M$ exponentially decaying terms. To show why this is true, we can write the Master equation for such a system as $\frac{d}{dt}\vec{P} = A\vec{P}$ where $\vec{P} \in \mathbb{R}^M$ is the vector of probabilities that the system is in each of the $M$ bound states, and $A$ is an $M \times M$ matrix of transition rates (including rates to the ruptured state). The solution to the Master equation is $P(t) = e^{At}\vec{P}_0 = \sum_{i=1}^{M} c_i \vec{v}_i e^{\lambda_i t}$, where $\vec{P}_0$ and the constants $c_i$ are set by the initial probabilities at $t = 0$ (which can be taken to be Boltzmann distributed according to the energies of the bound states), and where $\lambda_i$ and $\vec{v}_i$ are the eigenvalues and eigenvectors of $A$ (note that all $\lambda_i < 0$). Then, the probability that the bond remains unbroken at time $t$ is $P_{\text{unbroken}}(t) = \sum_j \vec{P}_j(t) = \sum_{i=1}^{M} c_i(\sum_j \vec{v}_{i,j})e^{\lambda_i t}$. The bond lifetime distribution is given by $p_\tau(t) = -\frac{d}{dt}P_{\text{unbroken}}(t) = \sum_{i=1}^{M} a_i e^{\lambda_i t}$, where $a_i = -c_i(\sum_j \vec{v}_{i,j})\lambda_i$. Therefore, as claimed, $p_\tau(t)$ has a multi-exponential form, with the number of exponential terms equal to the number of bound states.

### IV. TWO-STATE MODEL IN FIG. 3C

In Fig. 3C of the main text, we compare the bond lifetime distribution data to predictions of our model and of the two-state model. The two-state model [2] is defined by two bound states (states 1 and 2) and a ruptured state (state R), with dynamics given by the Master equation

$$\frac{d}{dt}\begin{bmatrix} P_1 \\ P_2 \end{bmatrix} = \begin{bmatrix} -k_{1R} - k_{12} & k_{21} \\ k_{12} & -k_{2R} - k_{21} \end{bmatrix} \begin{bmatrix} P_1 \\ P_2 \end{bmatrix} \tag{SM1}$$

where $k_{ij} = k_{ij}^o e^{\beta f \Delta x_{ij}}$ is the force-dependent transition rate from state $i$ to $j$. There are eight free parameters: $k_{ij}^o$ and $\Delta x_{ij}$ for $ij = 12, 21, 1R, 2R$.

The two-state model can describe slip-catch-slip bonding, yet it predicts a bond lifetime distribution that is inconsistent with data for E-selectin's slip-catch-slip behavior. To show this, we find the parameters $k_{ij}^o$ and $\Delta x_{ij}$ that give the best fit to the E-selectin mean lifetime data, then plot the predicted lifetime distribution using these parameters. Fig. SM2 shows the predicted mean lifetimes for the two-state model compared to data, with parameters listed in the figure caption. Fig. 3C of the main text shows the bond lifetime distribution with these parameters for $f = 48$ pN.

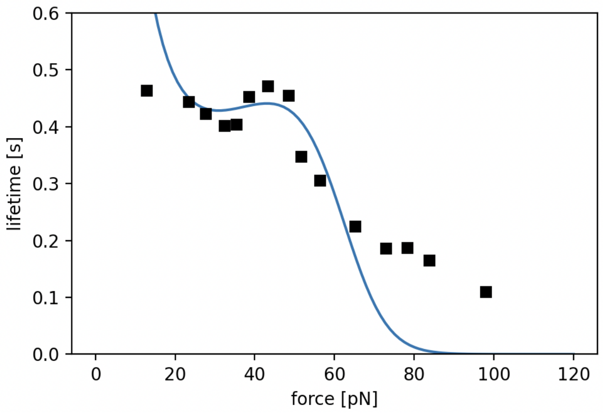


Fig. SM 2. Two state model (solid blue) compared to E-selectin mean lifetime data (black squares). Data from [3]. The best-fit parameters are: $k^o_{1R} = 1.07 \times 10^{-4}$ s$^{-1}$, $k^o_{2R} = 0.829$ s$^{-1}$, $k^o_{21} = 0.117$ s$^{-1}$, $k^o_{12} = 0.756$ s$^{-1}$, $\beta\Delta x_{1R} = 0.127$ pN$^{-1}$, $\beta\Delta x_{2R} = 0.0819$ pN$^{-1}$, $\beta\Delta x_{21} = -0.0219$ pN$^{-1}$, $\beta\Delta x_{12} = -8.05 \times 10^{-3}$ pN$^{-1}$.

### V. REMOVING HARMONIC CONTRIBUTIONS TO $V_f$ THROUGH COARSE-GRAINING

Harmonic contributions to a system's free energy can be removed by course-graining to produce a lower-dimensional free energy function with the same force-dependent energy barrier as the original free energy. Consider an $N$-dimensional free energy (or potential energy surface) of the form

$$U(\vec{Q}) = \tilde{U}(\boldsymbol{x}(\vec{Q})) + \frac{1}{2}\vec{Q}^T \Omega \vec{Q} - f\vec{\kappa} \cdot \vec{Q} \tag{SM2}$$

where $\vec{Q}, \vec{\kappa} \in \mathbb{R}^N$, and with $\boldsymbol{x} = P\vec{Q} \in \mathbb{R}^n$ with $n < N$, $P$ an $n \times N$ projection matrix, and $\Omega$ an $N \times N$ matrix. $\tilde{U}$ includes the non-harmonic contributions to the free energy.

This $N$-dimensional free energy can be reduced to an $n$-dimensional free energy of the form $V(\boldsymbol{x}) = \tilde{U}(\boldsymbol{x}) + \frac{1}{2}(\boldsymbol{x} - \boldsymbol{x}_0)^T A(\boldsymbol{x} - \boldsymbol{x}_0) - f\boldsymbol{\ell} \cdot \boldsymbol{x}$. This coarse-grained free energy has dimension equal to the dimension of $\tilde{U}$, the nonharmonic contribution.


[1] Wendy Thomas, Manu Forero, Olga Yakovenko, Lina Nilsson, Paolo Vicini, Evgeni Sokurenko, and Viola Vogel, "Catch-bond model derived from allostery explains force-activated bacterial adhesion," Biophysical journal **90**, 753–764 (2006).

[2] V Barsegov and D Thirumalai, "Dynamics of unbinding of cell adhesion molecules: transition from catch to slip bonds," Proceedings of the National Academy of Sciences **102**, 1835–1839 (2005).

[3] Annica M Wayman, Wei Chen, Rodger P McEver, and Cheng Zhu, "Triphasic force dependence of e-selectin/ligand dissociation governs cell rolling under flow," Biophysical journal **99**, 1166–1174 (2010).
